# Divergent Roles for cAMP–PKA Signaling in the Regulation of Filamentous Growth in *Saccharomyces cerevisiae* and *Saccharomyces bayanus*

**DOI:** 10.1101/311415

**Authors:** Ömur Kayikci, Paul M. Magwene

**Affiliations:** Department of Biology, Duke University, Durham, North Carolina, United States of America

**Keywords:** Gene network evolution, Signal transduction, cyclic-AMP, Fungal genetics

## Abstract

The cyclic AMP – Protein Kinase A (cAMP–PKA) pathway is an evolutionarily conserved eukaryotic signaling network that is essential for growth and development. In the fungi, cAMP–PKA signaling plays a critical role in regulating cellular physiology and morphological switches in response to nutrient availability. We undertook a comparative investigation of the role that cAMP-PKA signaling plays in the regulation of filamentous growth in two closely related budding yeast species, *Saccharomyces cerevisiae* and *Saccharomyces bayanus*. Using chemical and genetic perturbations of this pathway and its downstream targets we discovered divergent roles for cAMP-PKA signaling in the regulation of filamentous growth. While cAMP-PKA signaling is required for the filamentous growth response in both species, increasing or decreasing the activity of this pathway leads to drastically different phenotypic outcomes. In S. *cerevisiae*, cAMP-PKA inhibition ameliorates the filamentous growth response while hyper-activation of the pathway leads to increased filamentous growth; the same perturbations in S. *bayanus* result in the obverse. Divergence in the regulation of filamentous growth between S. *cerevisiae* and S. *bayanus* extends to downstream targets of PKA, including several kinases, transcription factors, and effector proteins. Our findings highlight the potential for significant evolutionary divergence in gene network function, even when the constituent parts of such networks are well conserved.

---

The cyclic AMP-Protein Kinase A (cAMP-PKA) pathway is an evolutionarily conserved signaling network that is important for the regulation of growth, differentiation, and development in animals, fungi, and amoebae (Toda et al. 1985; Zimmerman et al. 2015; D’Souza and Heitman 2001; Cho-Chung 2004; Das et al. 2007; Rinaldi et al. 2010; Gold et al. 2013; Loomis 2014). The basic principles of eukaryotic cAMP-PKA signaling are simple - in response to internal or external stimuli, increased adenylate cyclase activity causes a rise in intracellular cAMP levels. cAMP molecules bind to the regulatory domain of the PKA holoenzyme, releasing catalytic PKA subunits that phosphorylate downstream targets such as other kinases and transcription factors. cAMP production by adenylate cyclase is counter-balanced by cAMP breakdown via phosphodiesterases. Positive and negative feedback loops and temporally and spatially dynamic patterns further help to regulate cAMP-PKA activity (Toda et al. 1985, 1987; Belotti et al. 2012) In the model eukaryote, *Saccharomyces cerevisiae* (budding yeast), the cAMP-PKA signaling pathway helps to coordinate growth and cell fate decision-making in response to nutrient availability (Zaman et al. 2008; Gancedo 2013).

Filamentous growth is a cAMP-PKA regulated developmental response which is characterized by cell elongation, unipolar budding, physical attachment of mother and daughter cells, and increased adhesion to and invasion of growth substrates (Figure 1A). Nitrogen limitation is the primary trigger for filamentous growth in diploid cells, whereas haploid cells undergo filamentous differentiation in response to glucose limitation. The diploid filamentous growth response is also referred to as pseudohyphal growth, and we use both terms interchangeably in this study. *S. cerevisiae* filamentous differentiation is positively correlated with the activity of the cAMP-PKA pathway; genetic or biochemical manipulations that increase intracellular cAMP levels or PKA activity result in increased filamentous growth, while manipulations that decrease the net activity of the pathway ameliorate or abolish filamentous growth (Cullen and Sprague 2012; Gimeno and Fink 1994) (Figure 1B). Downstream targets of PKA include several transcription factors that regulate the expression of a cell wall glycoprotein, Flo11, required for filamentous growth in *S. cerevisiae* (Rupp et al. 1999; Lo and Dranginis 1998; Pan and Heitman 1999). Many of these same transcription factors are regulated in parallel by a MAP-kinase cascade (FG-MAPK). Both cAMP-PKA signaling and the FG-MAPK pathway are regulated by the Ras protein, Ras2.

**Figure 1.**
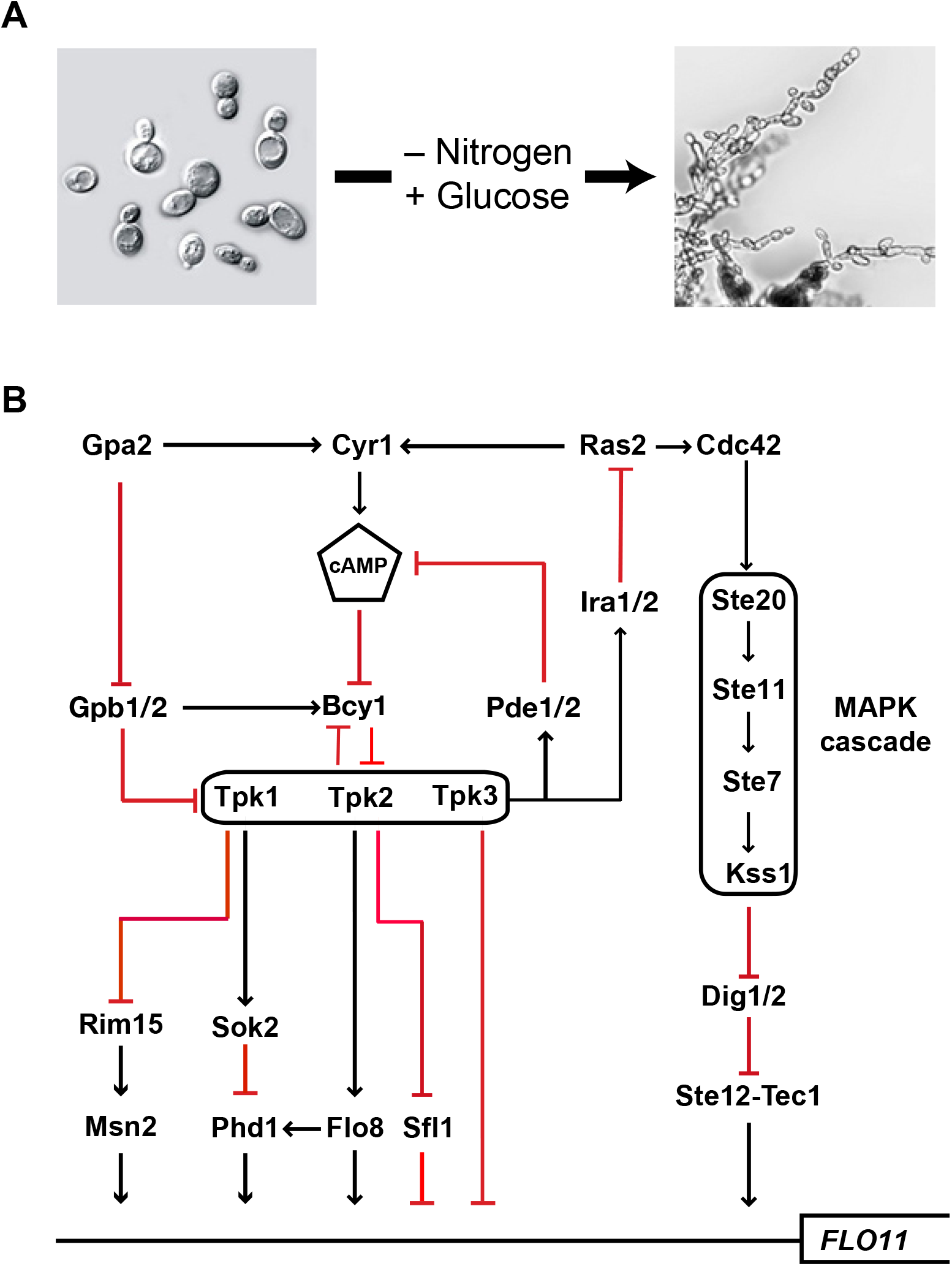
Filamentous growth in budding yeast. A) Upon nitrogen depletion, yeasts of the genus *Saccharomyces* undergo pseudohyphal differentiation in the presence of a fermentable carbon source, such as glucose. B) Flo11, a cell wall adhesin that is required for filamentous growth in *S. cerevisiae* is regulated in parallel by cAMP-PKA signaling and the filamentous growth MAP kinase pathway.

*S. cerevisiae* and related yeast within the *Saccharomyces sensu stricto* clade, provide a powerful comparative framework for understanding the evolution of gene networks (Cliften *et al*. 2003; Dujon 2010; Replansky et al. 2008; Hittinger 2013; Boynton and Greig 2014). Two additional species, *Saccharomyces paradoxus* and *Saccharomyces bayanus*, have received particular attention (Figure S1). *S. paradoxus*, the closest relative to *S. cerevisiae*, is primarily isolated from woodland areas and shows little genomic evidence of human facilitated admixture (Sampaio and Gonçalves 2008; Fay and Benavides 2005; Johnson *et al*. 2004; Naumov et al. 1998; Kowallik et al. 2015). *S. bayanus*, a lager yeast, is more distantly related to *S. cerevisiae* and *S. paradoxus*, and recent studies suggest that the phylogenetic history of the *S. bayanus* lineage involves a complex history of interspecific hybridization, facilitated by human activity (Sampaio and Gonçalves 2008; Naumov and Naumova 2011; Masneuf-Pomarède et al. 2010; Rodríguez et al. 2014; Pérez-Través *et al*. 2014). Since the nomenclature for the *S. bayanus* species complex is in flux (Hittinger 2013), for the purposes of this study we have adopted a conservative approach and refer to all strains belonging to this species complex as *S. bayanus*. *S. cerevisiae*, *S. paradoxus*, and *S. bayanus* display different physiologies, such as distinct differences in growth and survival strategies (Hittinger 2013; Borneman and Pretorius 2015). Within the *Saccharomyces* lineage, all of the major components of cAMP-PKA pathway are conserved.

In the present study we marshal phenotypic, biochemical, and genetic data to demonstrate that the regulation of filamentous growth by the cAMP-PKA signaling pathway has diverged significantly between *S. bayanus* and *S. cerevisiae*. We find that high levels of cAMP signaling have opposite effects on filamentous growth among these three species, promoting filamentous growth in both *S. cerevisiae* and *S. paradoxus* while inhibiting the filamentous response in *S. bayanus*. Divergent affects on the filamentous growth phenotype extend to downstream targets of PKA as well. In sum, our findings demonstrate that significant rewiring of the cAMP signaling pathway has occurred at multiple points in the cAMP-PKA gene network among the closely related species of the *Saccharomyces sensu stricto*. Our results, taken together with other recent findings regarding intraspecific variation and the potential for rapid evolution of cAMP-PKA signaling in response to selection, suggest that the cAMP-PKA pathway may be an evolutionary hot-spot for the accumulation of alleles that contribute to adaptation to novel nutrient niches.

## Materials and Methods

### Strains

Laboratory and environmental isolates of *S. cerevisiae, S. paradoxus*, and *S. bayanus*, and their corresponding pseudohyphal growth phenotypes are provided in Tables S1, S2, and S3. Mutants strains used in this study are given in Table S7. For *S. bayanus*, homozygous null mutants were generated in the NCYC365 background using KanMX4 deletion-cassette (Goldstein and McCusker 1999) with the standard PEG/LiAc protocol modified at the heat shock step, which was performed at 37rC for 45 minutes. The generated mutants, were confirmed with PCR and Sanger sequencing using primers listed in Table S8.

### Media and Phenotyping

Strains were grown overnight in YPD to a density of 2 ×10^7^ cells/ml. The cells were then washed twice in sterile water and 10^6^ cells were transferred to agar plates. Pseudohyphal growth was assayed using a modified SLAD medium (SLAD-1%) consisting of 0.17% YNB AA/AS, 1% dextrose, 50 M ammonium sulfate, and 2%Noble agar (Gimeno et al. 1992). For drug treatments, plates were supplemented with the indicated concentrations of cAMP (Enzo), 8-Bromoadenosine 3’,5’-cyclic monophosphate [8-Br-cAMP] (Sigma), 3-isobutyl-1-methylxanthine [IBMX] (Sigma), H-89 (Sigma), MDL 12,330A [MDL] (Sigma), and 2-5-Dideoxyadenosine [ddAdo] (Santa Cruz). For phenotyping, *S. cerevisiae* and *S. paradoxus* were incubated at 30řC, and *S. bayanus* strains were incubated at room temperature (RT). The strains were scored for pseudohyphal growth by the presence or absence of cellular projections at the colony edges, and the response was evaluated qualitatively as increased (+), decreased (−), or no change (Ø) relative to wild-type at 72 hours post plating. Images were collected using a Leica stereo microscope.

## Results

### Intra- and interspecific variation in pseudohyphal growth

We measured filamentous growth under nitrogen limitation in a genetically diverse panel of *S. cerevisiae* (36 strains), *S. paradoxus*(35 strains), and *S. bayanus* (36 strains) strains (Tables S1, S2, and S3). We adopted a binary classification system, rating each strain as pseudohyphal or non-pseudohyphal after 72 hours of growth on low-nitrogen growth medium (SLAD; see methods). Scoring was done via microscopic observation of the periphery of colonies for the presence of elongated cells, unipolar budding, and characteristic multicellular arrangements of cells into chains and branches. A similar fraction of strains in both *S. cerevisiae* and *S. bayanus* exhibited pseudohyphal growth (63.8% and 61.1% respectively). Only 31.4% of *S. paradoxus* strains showed pseudohyphal after 72 hours of growth on SLAD. For all three species, there was significant variation in the strength of the pseudohyphal response among those strains capable of filamentous growth.

### Exogenous cAMP inhibits pseudohyphal growth in S. bayanus

Previous studies have demonstrated that application of exogenous cAMP to the growth medium increases the propensity to form pseudohyphae in *S. cerevisiae*, and can restore pseudohyphal growth in mutants with reduced cAMP production (Lorenz and Heitman 1997; Kübler et al. 1997). This effect presumably mimics the increased activity of the endogenous adenylate cyclase. To test the generality of this effect across the *Saccharomyces sensu stricto* clade, we grew pseudohyphal and a non-pseudohyphal strains of *S. cerevisiae, S. bayanus*, and *S. paradoxus* under nitrogen-limiting conditions with various concentrations of exogenous cAMP (1 mM, 3 mM, 10 mM) added to the growth media. Most non-pseudohyphal *S. cerevisiae* and *S. paradoxus* isolates displayed a strong pseudohyphal phenotype in response to the presence of cAMP, exhibiting numerous filamentous extensions at the colony perimeter as well as increased invasiveness. Similarly, strains of *S. cerevisiae* and *S. paradoxus* that already exhibited the ability to undergo pseudohyphal growth showed a qualitative increase in the response upon cAMP treatment. In striking contrast, exogenous cAMP treatment was ineffective in inducing pseudohyphal differentiation in *S. bayanus* strains. Not only was the cAMP treatment ineffective in inducing the response in non-pseudohyphal *S. bayanus*isolates but, surprisingly, cAMP treatment suppressed filamentous differentiation in more than half of the normally pseudohyphal *S. bayanus* strains (Figure 2 and Table S4). We also tested the effect of the cAMP analog 8-Br-cAMP, which is reported to be more membrane permeant and resistant to degredation by phosphodiesterases (Schaap et al. 1993). 8-Br-cAMP at a concentration of 500 M produced a reduction of pseudohyphal growth in *S. bayanus* and an increase in *S. cerevisiae* comparable to approximately 3 mM cAMP (Figure S4).

**Figure 2.**
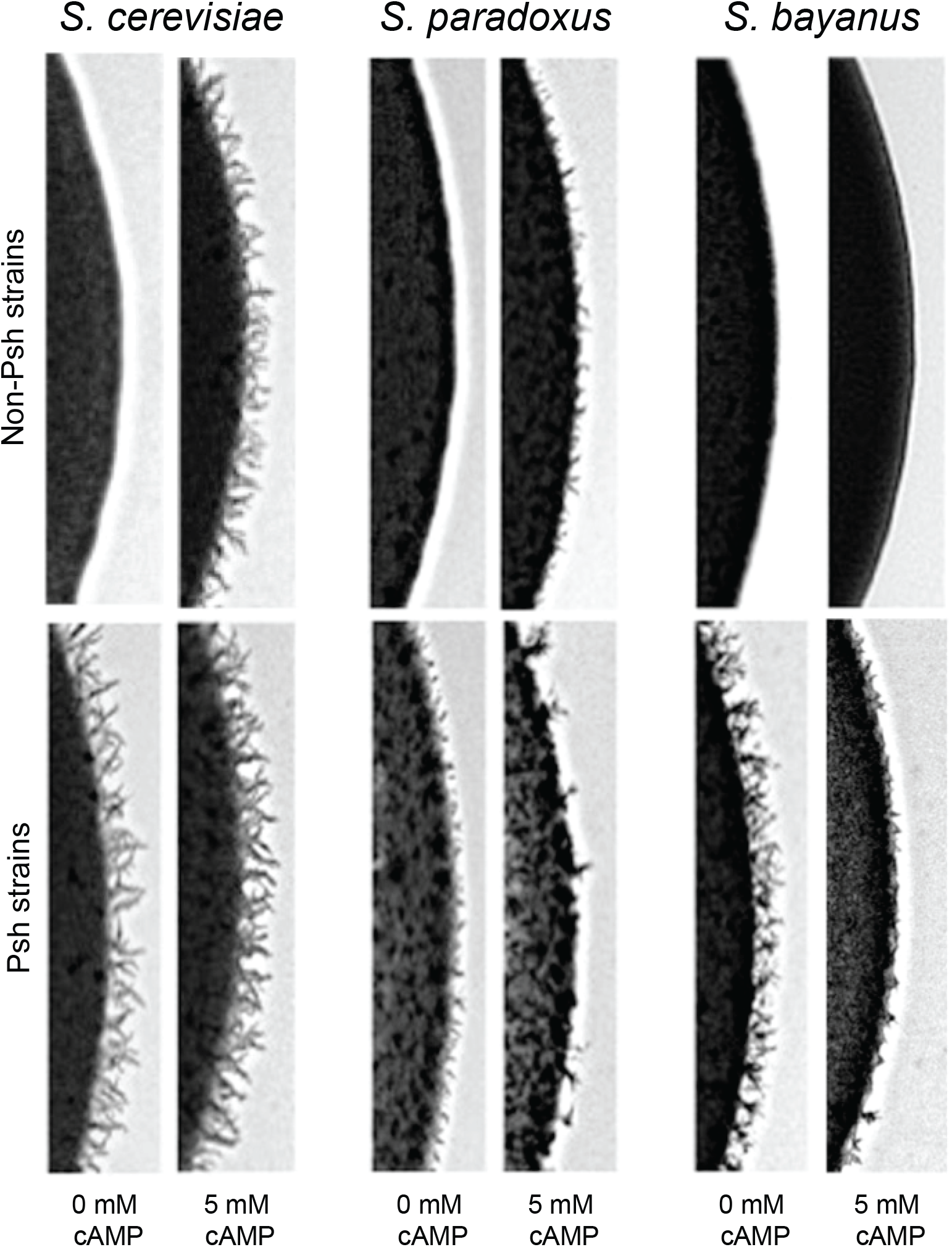
Exogenous cAMP inhibits pseudohyphal growth in *S. bayanus*. Pseudohyphal phenotypes are shown for a pseudohyphal (Psh) and a non-pseudohyphal (Non-Psh) strain of each species, grown in the presence of 5 mM cAMP. cAMP treatment promotes pseudohyphal growth in *S. cerevisiae* and *S. paradoxus* but inhibits pseudohyphal growth in *S. bayanus*.

### Chemical manipulation of the cAMP-PKA Pathway

In order to further explore the surprising effect that exogenous cAMP had on filamentous growth in *S. bayanus*, we scored pseudohyphal growth in the presence of four additional chemical agents that have been shown to modify the activity of key enzymes involved in cAMP-PKA signaling. MDL-12,330A and 2-5 Dideoxyadenosine (ddAdo) directly inhibit the activity of adenylate cyclase (Cutuli et al. 2000; Guellaen et al. 1977), and thus should decrease intracellular cAMP levels. H89 is a protein kinase A inhibitor with broad specificity (Murray 2008), but is likely to decrease PKA activity. IBMX is an inhibitor of phosphodiesterases (Van Lookeren Campagne et al. 1990), and thus would tend to favor accumulation of cAMP in cells. Treatment with both MDL and ddAdo lead to a drastic decrease in pseudohyphal growth in *S. cerevisiae* and a modest decrease in *S. paradoxus*, but the filamentous response in *S. bayanus* in the presence of these agents is comparable to the untreated control (Figure S3). A 1 mM IBMX treatment increased the pseudohyphal response in both *S. cerevisiae* and *S. paradoxus*, while decreasing the density of pseuodohyphal projections on the margin of colonies in *S. bayanus* (Figure S4). A higher concentration of IBMX (3 mM), however, led to a dimunution of the response in all three species. The PKA inhibitor H-89 (50 M) had no discernible effects on pseudohyphal growth in *S. bayanus*, however there was a modest to complete loss of pseudohyphal growth in response to H89 in both *S. cerevisiae* and *S. paradoxus* (Figure S4).

Since *S. cerevisiae* and *S. bayanus* showed the greatest divergence of filamentous phenotypes in response to nutrient limitation and chemical manipulation, we chose to concentrate further investigations on these two species.

### MAPK functions similarly in S. cerevisiae and S. bayanus pseudohyphal response

Both the cAMP-PKA pathway and the FG-MAPK cascade are capable of inducing pseudohyphal growth in *S. cerevisiae*. To rule out differences in the contribution of the FG-MAPK cascade to filamentous growth in the two species, we carried out gene deletion experiments in *S. bayanus* to confirm that FG-MAPK mutant phenotypes are similar to those previously reported for *S. cerevisiae*. Using drug resistance markers, we created deletion mutants of *STE7, STE12, TEC1*, and *DIG1*. The mutants of the positively contributing MAPK components, *ste7*, *ste12*, and *tec1*, exhibited smooth colony edges and a lack of invasiveness. The deletion of the negative element, *DIG1*, led to an increase in the filamentous response (Figure S2). These results are consistent with phenotypes observed for the same mutants in *S. cerevisiae* (Cook et al. 1996; Madhani and Fink 1997; Oehlen and Cross 1998; Roberts and Fink 1994).

### The cAMP-PKA pathway is required for the filamentous response in both S. cerevisiae and S. bayanus

Having ruled out the FG-MAPK pathway as a likely candidate for the differences observed between *S. cerevisiae* and *S. bayanus*, we proceeded with systematic genetic manipulation of key genes in the cAMP-PKA pathway. We deleted 11 genes encoding elements of the cAMP pathway in *S. bayanus*, and compared the resulting filamentous growth phenotypes to those of the same mutants in *S. cerevisiae*. Unlike FG-MAPK mutants, we found that the effects of gene deletions in the cAMP-PKA pathway often differed in terms of observed phenotypes between *S. bayanus* and *S. cerevisiae*. We classified our observations into two categories of effects: 1) mutants with similar phenotypes and 2) mutants with opposite effects (Table S5).

The first category of mutants, exhibiting similar phenotypes in both species, included *gpa2, tpk1, tpk2*, and *tpk3*. Deletion of *TPK2* ameliorates the FG response in both *S. bayanus* and *S. cerevisiae*, indicating that this PKA subunit is required for induction of filamentous growth in both species (Figure 3) (Robertson and Fink 1998; Pan and Heitman 1999). *tpk1* and *tpk3* mutants have the opposite effect relative to *tpk2*, showing increased pseudohyphal growth in *S. bayanus* as has been previously reported for *S. cerevisiae* (Robertson and Fink 1998; Pan and Heitman 1999). This confirms that the distinct roles of the PKA subunits in the regulation of filamentous growth is conserved between the two species. Gpa2 is an activator of the adenylate cyclase Cyr1, and an inhibitor of the kelch repeat proteins Gpb1 and Gpb2. The *gpa2* mutants show a loss of pseudohyphal growth in both species (Figure 4). The *gpb1* and *gpb2* mutants in *S. bayanus*show a slight increase in pseudohyphal growth (Figure S6), similar to what has been reported for *S. cerevisiae* (Harashima and Heitman2002).

**Figure 3.**
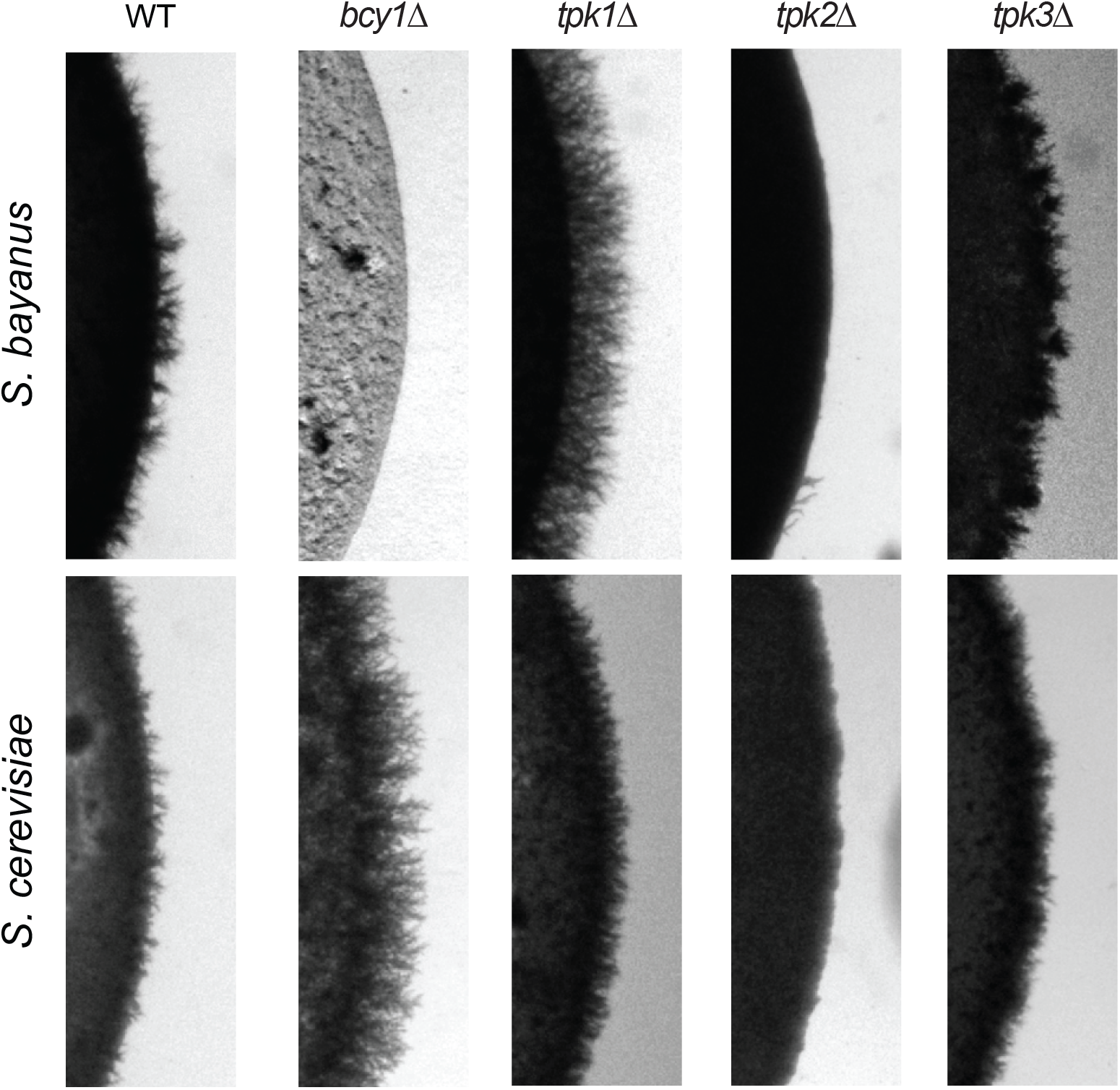
Mutations of subunits of the PKA holoenzyme have both similar and opposite effects on filamentous growth in *S. cerevisiae* and *S. bayanus*. The catalytic subunit *TPK2* promotes pseudohyphal growth in both species, while *TPK1* and *TPK3* are negative regulators of pseudohyphal growth. Deletion of the PKA catalytic subunits leads to parallel phenotypes in the two species. By contrast, deletion of the regulatory subunit, *BCY1*, results in hyper-filamentous growth in *S. cerevisiae*, but extremely slow growth with no pseudohyphae in *S. bayanus* (see also supplementary Figure S5). Mutants are on 1278b and NCYC 365 backgrounds for *S. cerevisiae* and *S. bayanus*, respectively.

**Figure 4.**
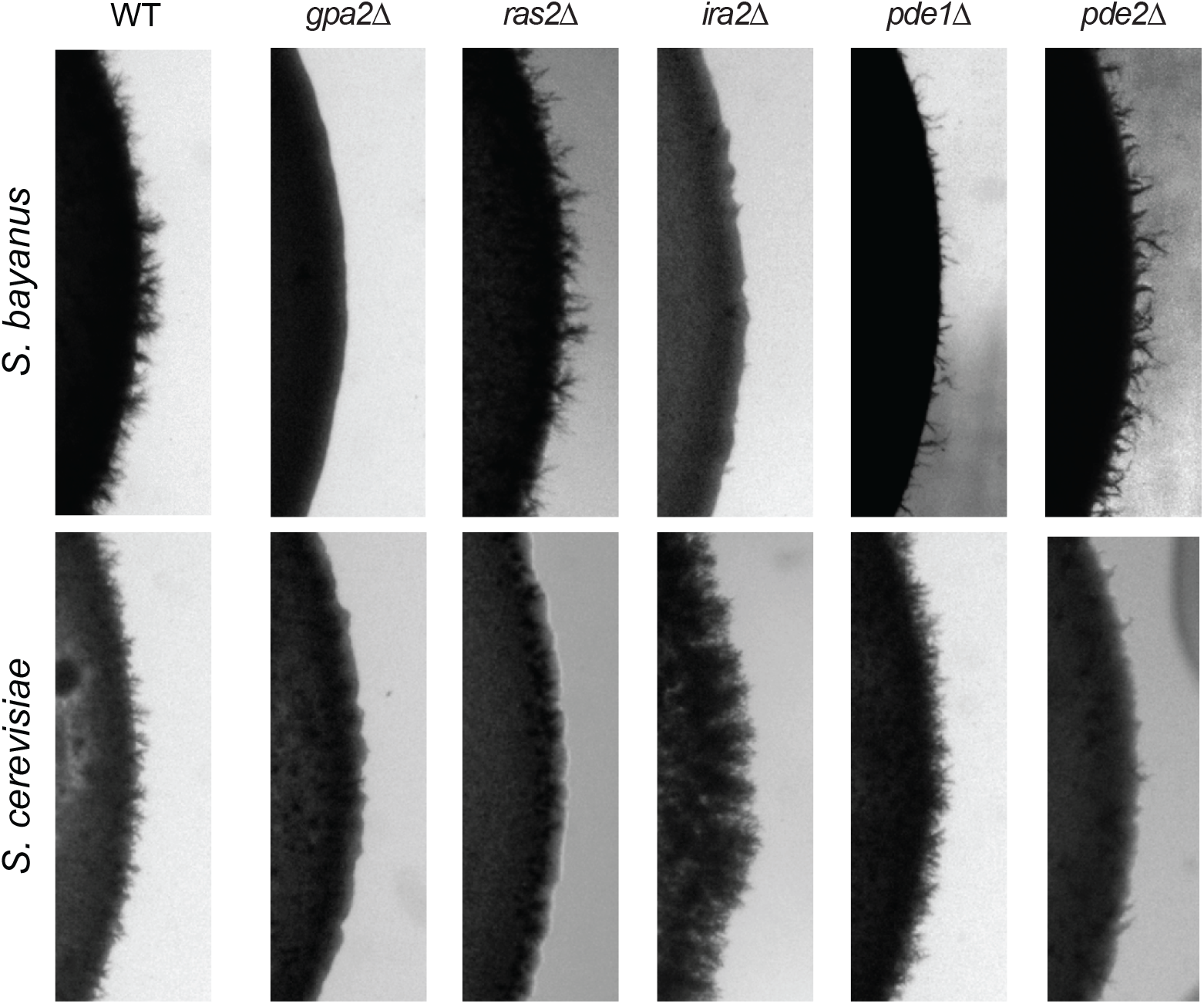
Mutations that affect cAMP levels have primarily opposite effects on filamentous growth in *S. cerevisiae* and *S. bayanus*. With the exception of *gpa2*, deletion mutations that affect adenylate cyclase activity or cAMP concentration show opposite phenotypic effects in *S. cerevisiae* and *S. bayanus*. See text for further discussion.

Mutants with opposite phenotypes in the two species included *ras2, pde1, pde2, ira2*, and *bcy1* (Figures 3 and 4 and summarized in Table S5). The *ras2* mutants show a strong decrease of filamentous growth in *S. cerevisiae*, but no decrease in *S. bayanus*. The *ira2* mutants show an increase of filamentous growth in *S. cerevisiae*, and a strong decrease in *S. bayanus*. The *pde1* mutants show an increase in filamentous growth in *S. cerevisiae*, and a strong decrease in *S. bayanus*, while *pde2* mutants show a decrease of filamentous growth in *S. cerevisiae* and no change or a slight increase in *S. bayanus. bcy1* mutants in *S. cerevisiae* showed abundant pseudohyphae, while the same mutant in *S. bayanus* is very slow growing and shows insufficient growth after 72 hours to score FG. However, if *S. bayanus bcy1* mutants are allowed to grow for 10 days they eventually form a colony, but show no pseudohyphae (Figure S5).

### Mutant phenotypes of targets of PKA in S. bayanus

We next examined the phenotypic effects of knockout mutants of four transcription factors – Flo8, Phd1, Sfl1, and Msn2 – that are targets of PKA, and which are known to play key roles in regulating pseudohyphal growth in *S. cerevisiae* (Figure 5). Flo8 and Phd1 are positive regulators of the pseudohyphal response; while Sfl1 is a repressor. All three are thought to modify pseudohyphal growth primarily through transcriptional regulation of *FLO11* (see below). Msn2 is a stress responsive transcription factor that is regulated by both PKA and the TOR pathway.

**Figure 5.**
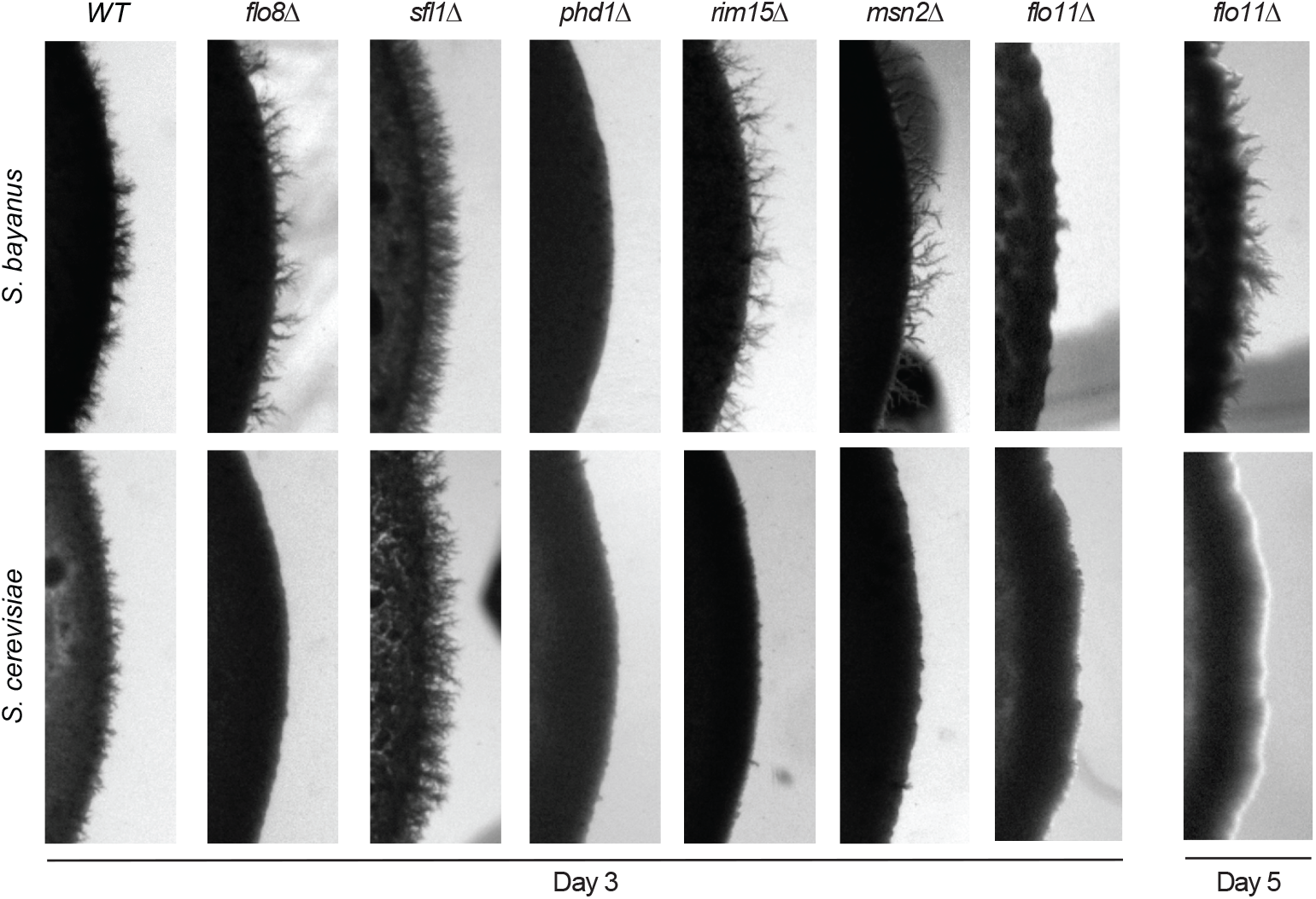
Downstream targets of cAMP-PKA signaling show a mix of similar and divergent pseudohyphal responses. Pseudohyphal phenotypes of*flo8* mutations differ between *S. cerevisiae* and *S. bayanus*, but the response upon deletion of *SFL1* and *PHD1* is conserved. Deletion of *FLO11* eliminates pseudohyphal growth completely in *S. cerevisiae;* in contrast, deletion of *FLO11*in *S. bayanus* causes a delay in the pseudohyphal response. At day three of observation filamentous growth is absent in the *flo11* mutants of both *S. cerevisiae* and *S. bayanus*, but *S. bayanus flo11* mutants start to exhibit pseudohyphal projections by day five.

The *phd1* and *sfl1* mutant phenotypes are identical between *S. cerevisiae* and *S. bayanus*, with *PHD1* deletion mutants showing a loss in filamentous growth and *SFL1* deletion mutants showing an increase in filamentous growth. Surprisingly, *flo8*mutants show opposite phenotypes in the two species, with complete abrogation of the pseudohyphal response observed in *S. cerevisiae* but no change in filamentous growth in *S. bayanus*. The *msn2* mutants show no loss of pseudohyphal growth in *S. bayanus*, while there is a complete loss of the phenotype in *S. cerevisiae*. Rim15 is a kinase that is a target of both PKA and TOR signaling, and in turn contributes to the regulation of Msn2. *rim15* mutants in *S. cerevisiae* show a loss of pseudohyphal growth, while *S. bayanus* mutants show comparable filamentous growth to the wild type background (Figure 5).

### FLO11 is not required for filamentous growth in S. bayanus

The cell wall glycoprotein Flo11 is regarded as one of the primary molecular effectors of pseudohyphal growth in *S. cerevisiae. flo11* mutants not only show a loss of pseudohyphal growth in *S. cerevisiae*, but also show an inability to form biofilms and complex colonies (Granek and Magwene2010; Granek et al. 2013; Zara et al. 2009). We compared *flo11* mutants in both *S. cerevisiae* and *S. bayanus* over the course of five days. *S. cerevisiae flo11* mutants show no sign of pseudohyphal growth, even up to five days post-plating. *S. bayanus* mutants show little filamentous growth at day three post-plating, but begin to show pseudohyphae at the colony margin at day 4, and show substantial pseudohyphae by day 5 (though less than WT) (Figure 5). We conclude that *flo11* deletion delays the expression of filamentous growth in *S. bayanus*, and thus may be a key effector of in both species, but our finding also suggests a Flo11 independent mechanism for producing pseudohyphae in *S. bayanus*.

## Discussion

The findings we describe above, regarding the role of cAMP-PKA signaling in the regulation of pseudohyphal growth in *S. bayanus*, are surprising in a number of respects. In *S. cerevisiae*, cAMP-PKA signaling plays an unambiguously positive role in the regulation of filamentous growth. Chemical and genetic manipulations that increase cAMP-PKA signaling lead to increased filamentous growth in *S. cerevisiae*, while perturbations that decrease cAMP-PKA signaling reduce the strength of the pseudohyphal response. In contrast, we find that in *S. bayanus*, perturbations that are predicted to increase intracellular levels of cAMP lead to a decrease in the filamentous growth response. These differences between the two species exist despite the fact that the core elements of the cAMP-PKA signaling network are highly conserved at the sequence level throughout the *Saccharomyces sensu stricto* species (Table S6).

### Chemical and genetic manipulation of cAMP levels produces divergent phenotypes in S. cerevisiae and S. bayanus

The application of exogenous cAMP exaggerates the pseudohyphal response in *S. cerevisiae* and *S. paradoxus*, but attenuates the pseudohyphal switch in *S. bayanus* (Figure 2). Pharmacological agents that modulate cAMP levels also show contrasting effects between *S. cerevisiae* and *S. bayanus* (Figures S3 and S4). Consistent with the results by chemical manipulation, genetic perturbation of the feedback mechanisms controlling cAMP levels results in starkly contrasting phenotypes between *S. cerevisiae* and *S. bayanus*. For example, knockouts of *PDE1*and *IRA2* increase intracellular cAMP levels and as a consequence *pde1* and *ira2* mutants exhibit exaggerated pseudohyphal growth in *S. cerevisiae* (Cullen and Sprague2012). The same mutations in *S. bayanus*, lead to a striking reduction in pseudohyphal growth. *RAS2* mutants, which show a loss of pseudohyphal growth in *S. cerevisiae*, have wild type pseudohyphal phenotypes in *S. bayanus*. The one exception to the pattern is the phenotypes observed for *gpa2* mutants, where both *S. cerevisiae* and *S. bayanus* show a loss of pseudohyphal growth.

### PKA mutations and downstream targets produce a mixture of similar and dissimilar phenotypes

In contrast to the generally divergent phenotypes exhibited by *S. cerevisiae* and *S. bayanus* upon manipulation of cAMP levels, the results we observed for mutants and chemical agents that affect PKA activity showed a mixture of similar and divergent phenotypes between the two species. Deletions of the PKA regulatory subunit, *BCY1*, which inhibits PKA activity, shows strong differences between the species. *bcy1* mutants show hyper filamentous growth in *S. cerevisiae*, while the same mutant is slow-growing and non-pseudohyphal in *S. bayanus*. However, deletions of the PKA catalytic subunits Tpk1, Tpk2, and Tpk3 produced identical phenotypes in both *S. cerevisiae*and *S. bayanus*, with *tpk1* and *tpk3* mutants both showing increased pseduohyphal growth while *tpk2* mutants show decreased pseudohyphal growth.

At the level of downstream targets of PKA, we again see a mix of similar and divergent phenotypes between *S. cerevisiae*and *S. bayanus* among deletion mutants. The transcription factors Phd1 and Sfl1 play similar roles in both species, however deletions of the transcription factors Flo8 and Msn2 produced opposite responses when comparing the species. The ability of *S. bayanus* to produce pseudohyphae in the absence of Flo8p is especially surprising as this deletion completely abrogates pseudohyphal growth in*S. cerevisiae* (Liu et al. 1996).

### Flo11 is partially dispensable for pseudohyphal growth in S. bayanus

In *S. cerevisiae* both the cAMP-PKA pathway and the filamentous growth MAPK pathway jointly regulate FLO11, a cell wall adhesin that is thought to be critical for nutrient-induced pseudohyphal growth. Loss-of-function or deletion mutations of *FLO11* eliminate nutrient-induced pseudohyphal growth in *S. cerevisiae* (Cullen and Sprague 2012). As we describe above, *S. bayanus flo11* mutants are slow to manifest pseudohyphal growth, but do eventually exhibit pseudohyphae, though the strength of the pseudohyphal response is reduced relative to wild-type. *FLO11* independent regulation of filamentous growth is not totally without precedent. For example, Lorenz et al. (Lorenz et al. 2000) reported that *FLO11* is dispensable for pseudohyphal growth in the presence of 1% butanol and Halme et al. (Halme et al. 2004) found that ira1 flo11 mutants can undergo FLO10 dependent pseudohyphal growth.

### The FG-MAPK cascade is conserved between S. cerevisiae and S. bayanus

In contrast to the numerous differences we documented with respect to the cAMP-PKA pathway, the genetic effects of perturbations to the filamentous growth MAPK cascade appears to be conserved between *S. cerevisiae* and *S. bayanus*, with both species showing similar mutant phenotypes for all the genes tested in this pathway. This conservation of genetic effects for FG-MAPK mutants holds even though previous studies have demonstrated significant divergence between *S. cerevisiae* and *S. bayanus* in the genes regulated by Ste12 and Tec1, two transcription factors that are targets of the FG-MAPK pathway and which contribute to the regulation of pseudohyphal growth (Borneman et al. 2007; Martin et al. 2012).

### Speculative Model and Future Directions

How might we integrate the findings presented above into a model for the role that cAMP-PKA signaling plays in the regulation of pseudohyphal in *S. bayanus?* Two broad patterns emerge from our chemical and genetic perturbations. The first is that some level of PKA activity is required for pseudohyphal growth in both *S. cerevisiae* and *S. bayanus*. The second is that high levels of cAMP are inhibitory of pseudohyphal growth in *S. bayanus*, while promoting pseudohyphal growth in *S. cerevisiae*.

Particularly interesting in this regard is the role of Bcy1, the PKA regulatory subunit that directly interacts with cAMP and hence is the critical mediator between intracellular cAMP levels and the downstream effects of PKA activity. High levels of cAMP relieve the inhibitory effects of Bcy1 on the PKA catalytic subunits – Tpk1, Tpk2, and Tpk3. Genetically, Tpk1 and Tpk3 are inhibitors of pseudohyphal growth while Tpk2 is an activator of pseudohyphal growth, as has been previously shown for *S. cerevisiae* (Robertson and Fink 1998; Pan and Heitman 1999), and as we show here for *S. bayanus*.

We hypothesize that *S. cerevisiae* and *S. bayanus* differ in the relative amount or activity of the PKA catalytic subunits, in response to changes in intracellular cAMP levels. Species specific differences in the relative expression of the different Tpk subunits, or their relative affinity for the PKA regulatory subunit, Bcy1, could favor a shift in the balance between Tpk1/Tpk3 versus Tpk2. We hypothesize that in *S. cerevisiae*, increased cAMP signaling favors greater activity of Tpk2, while in *S. bayanus* similar increases in cAMP favor greater Tpk1 and/or Tpk3 activity (Figure 6). This hypothesis can be tested in future studies using a combination of gene deletions and heterologous expression of the various PKA regulatory and catalytic subunits individually and in combination in both *S. cerevisiae* and *S. bayanus*.

**Figure 6.**
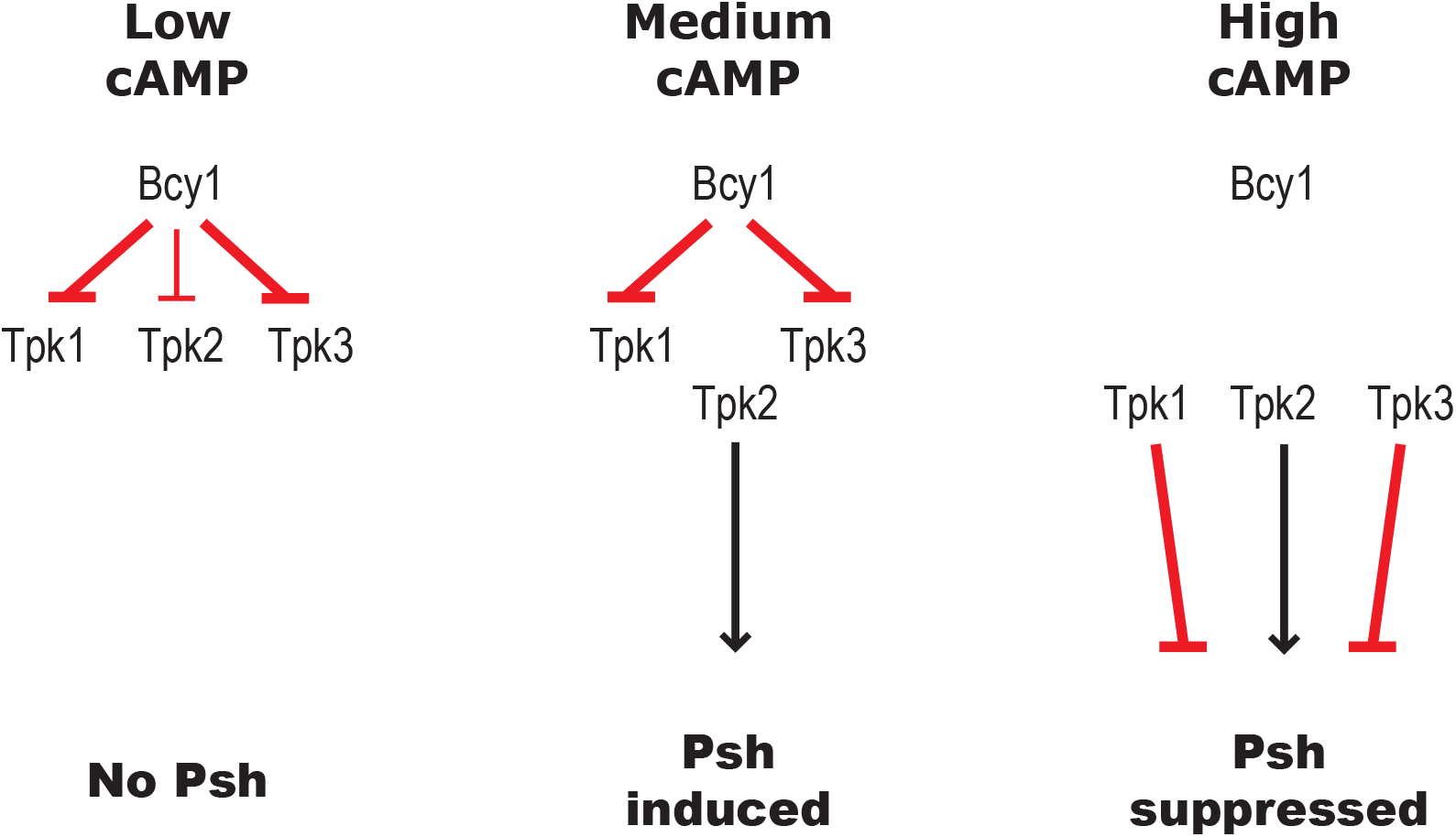
A proposed model for cAMP-PKA signaling in *S. bayanus*. To explain the differences in the regulation of pseudohyphal growth in *S. bayanus* and *S. cerevisiae*, we propose a model based on the relative strength of interactions (indicated by line weight) between the regulatory (Bcy1) and catalytic (Tpk1, Tpk2, Tpk3) PKA subunits. We hypothesize that moderate levels of cAMP signaling lead to the preferential release of the catalytic subunit Tpk2, a positive regulator of filamentous growth. At high concentrations of cAMP, the Tpk1 and Tpk3 (repressors of filamentous growth), are also released from the PKA holoenzyme, counteracting the effects of Tpk2 and suppressing pseudohyphal growth.

Our findings also point to differences in the relative importance of downstream effectors of PKA, particularly key transcription factors such as Msn2 and Flo8, for the regulation of pseudohyphal growth. This suggests that rewiring at the level of gene regulation also contributes to the differences between *S. cerevisiae* and *S. bayanus*.

More broadly we speculate that the differences we observe in the regulation of pseudohyphal growth by the cAMP-PKA pathway reflects physiological differences between the two species, not only with respect to nitrogen utilization, but other stresses as well (Blein-Nicolas et al. 2013; Masneuf-Pomarède *et al*. 2010; Serra *et al*. 2005).

### The cAMP-PKA pathway is an evolutionary hot-spot for adaptation in yeast

A number of other recent studies, focusing on variation *within S. cerevisiae*, highlight how standing genetic variation and de novo mutations in the cAMP-PKA pathway contribute to the genetic architecture of complex traits and adaptation to novel environments. These studies indicate that: 1) among environmental isolates of *S. cerevisiae* there is substantial genetic variation in the cAMP-PKA pathway and this variation affects a diversity of phenotypic traits (Granek et al. 2013; Taylor *et al*. 2016; Yadav et al. 2015); and 2) mutations that affect cAMP-PKA signaling are often among the earliest genotypic changes that are favored when yeast populations are subjected to selection in novel nutrient environments (Hong and Gresham2014; Li et al. 2018; Sato et al. 2016; Venkataram et al. 2016). Our findings, taken together with this growing body of work, thus point to the cAMP-PKA pathway as a major driver of evolutionary change in the the *Saccharomyces sensu stricto* species complex. Given the central role that cAMP-PKA signaling plays in the regulation of morphogenesis across the fungi (Hicks and Heitman 2007; Klengel et al. 2005; Dürrenberger et al. 1998), we expect that the central importance of this pathway for adaptation and evolution is likely to be recapitulated in many other fungal clades.

## Conclusions

This study highlights the evolutionary lability of the cAMP-PKA pathway among the species of the *Saccharomyces sensu stricto* complex. cAMP-PKA signaling is an key regulator of morphogenetic switches in response to environmental cues for the fungi generally (Boyce and Andrianopoulos 2015; Pérez Martín, José and Di Pietro, Antonio 2012; Turrà et al. 2014) and both inter- and intraspecific variation in cAMP-PKA signaling is likely to be an important genetic determinant of phenotypic variation in many fungal systems. More generally our findings exemplify the potential for conserved eukaryotic signaling pathways to diverge in the regulation of cellular phenotypes even among relatively closely related species.

## Acknowledgments

This research was funded National Science Foundation grant MCB 1330545. We thank John McCusker, Joseph Heitman, and Timothy Galitski for providing strains. We thank Andrew Alspaugh, Fred Nijhout, Amy Schmid, and Rytas Vilgalys for feedback and recommendations during the course of this study. We thank Debra Murray for proofreading and comments on the manuscript.

## Author Contributions

Conceived and designed the experiments: OK PM. Performed the experiments: OK. Analyzed the data: OK PMM. Wrote the paper: OK PMM.

## Supplementary Figures

**Figure S1.**
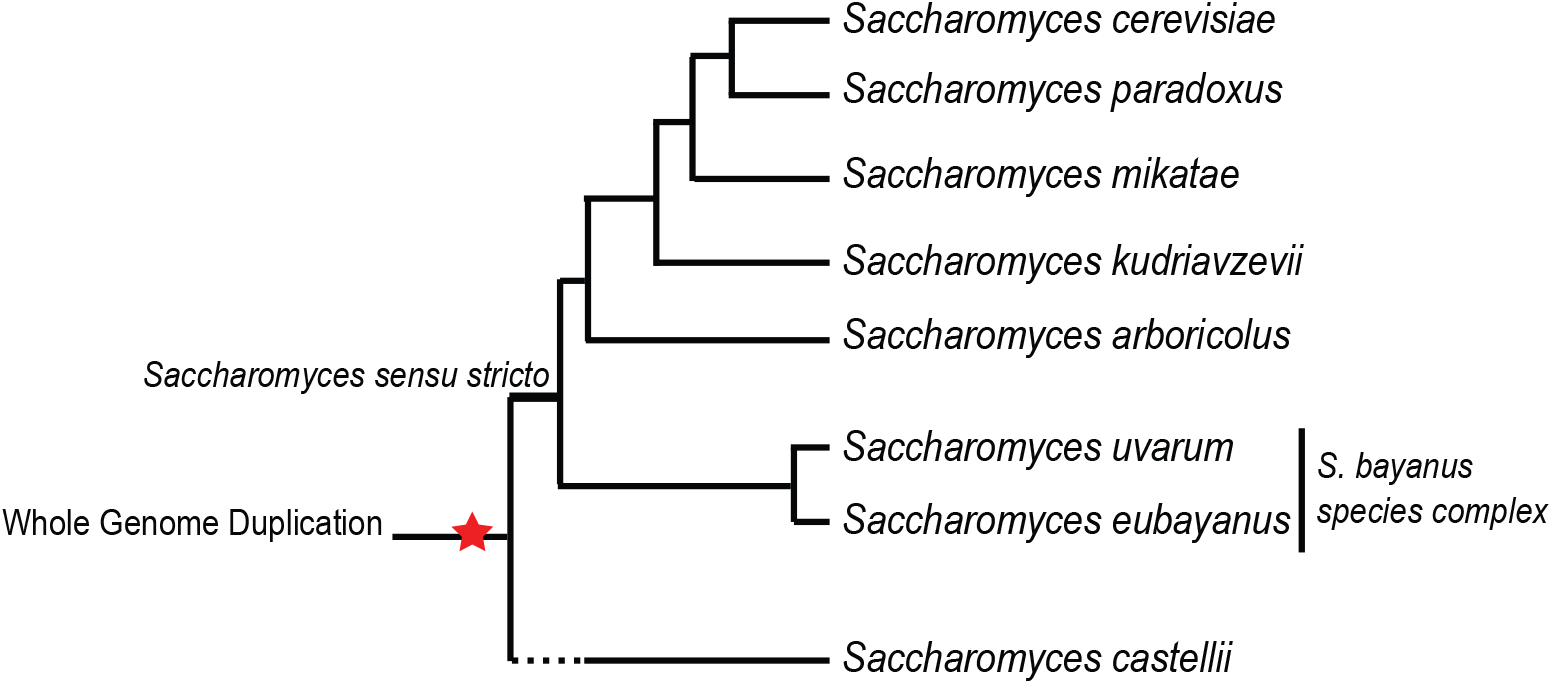
Closely related *Saccaromyces* species. *Saccharomyces* species inhabit a broad range of environments and exhibit different physiologies despite the short estimated-divergence time between lineages (5-20 mya).

**Figure S2.**
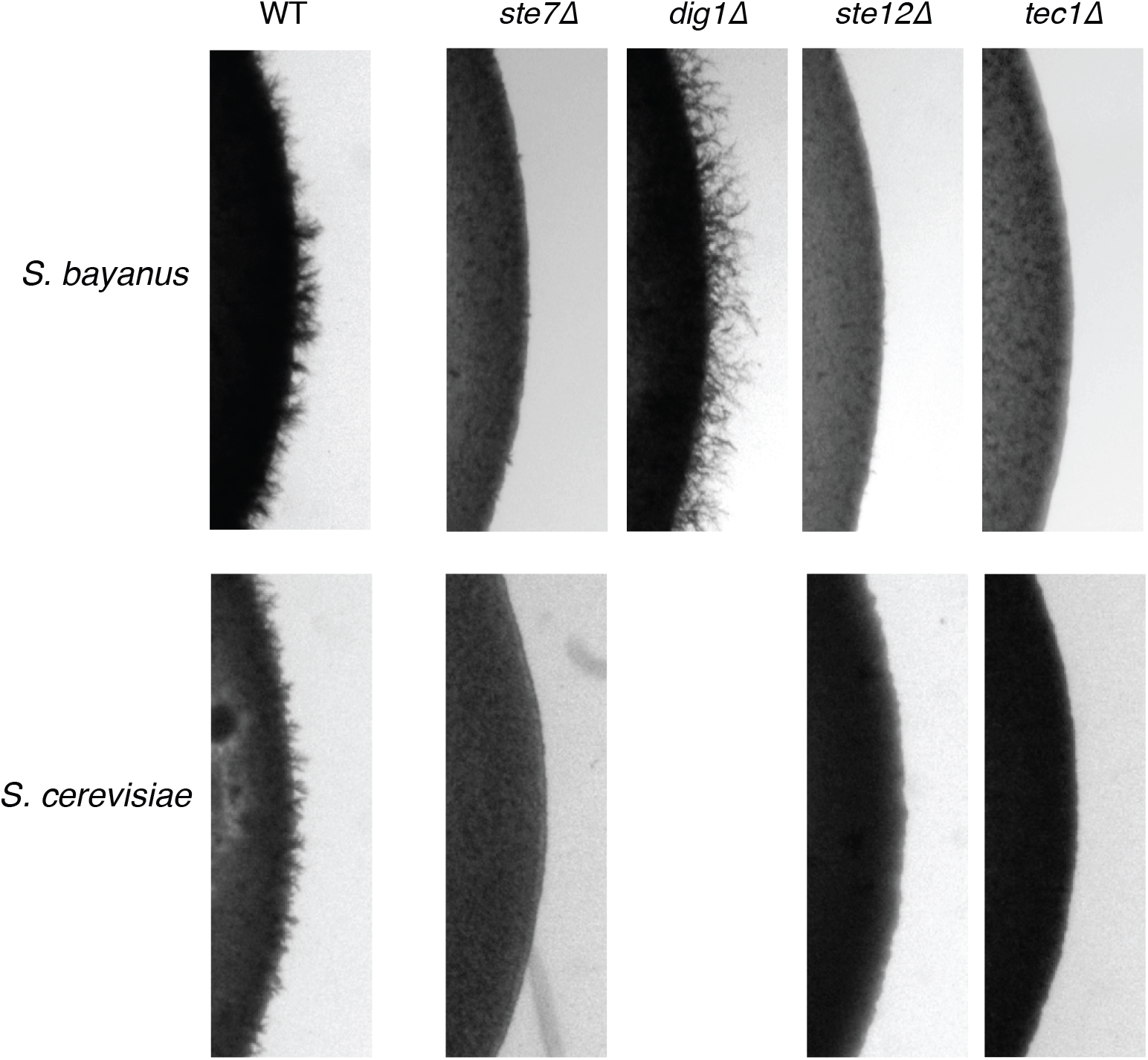
Disruption of MAPK signaling eliminates pseudohyphal growth in *S. cerevisiae* and *S. bayanus*, indicating that the cascade regulates the response positively in the two species. Mutants are in the 1278b and NCYC365 backgrounds for *S. cerevisiae* and *S. bayanus*, respectively.

**Figure S3.**
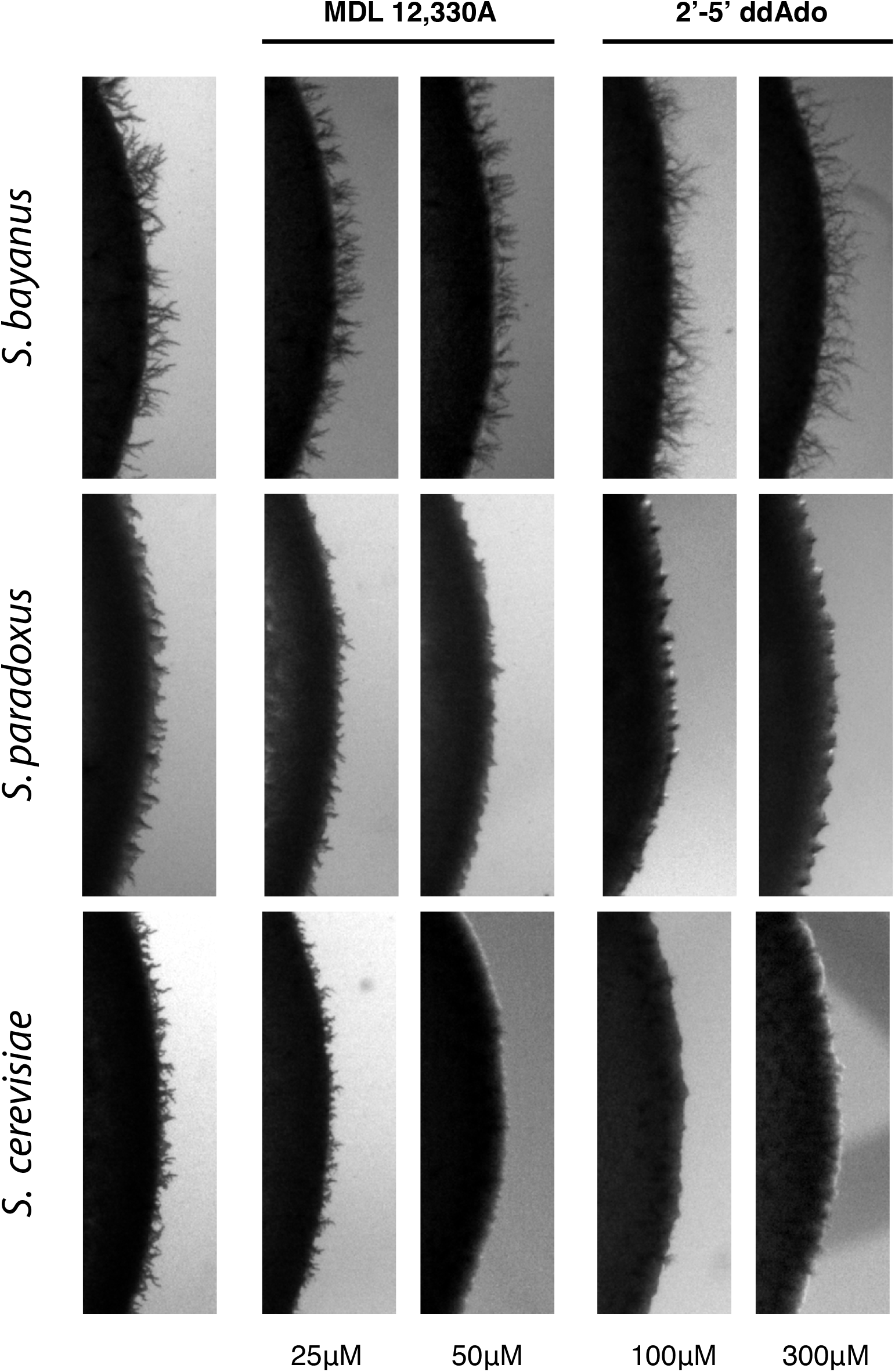
Pharmacological inhibition of cAMP synthesis and has divergent effect on pseudohyphal growth in *S. cerevisiae* and *S. bayanus*. MDL-12,330A and 2-5 Dideoxyadenosine inhibit adenylate cyclase activity in different ways. MDL-12,330A prevents the membrane localization of adenylate cyclase; 2-5 Dideoxyadenosine blocks the catalytic domain of adenylate cyclase.

**Figure S4.**
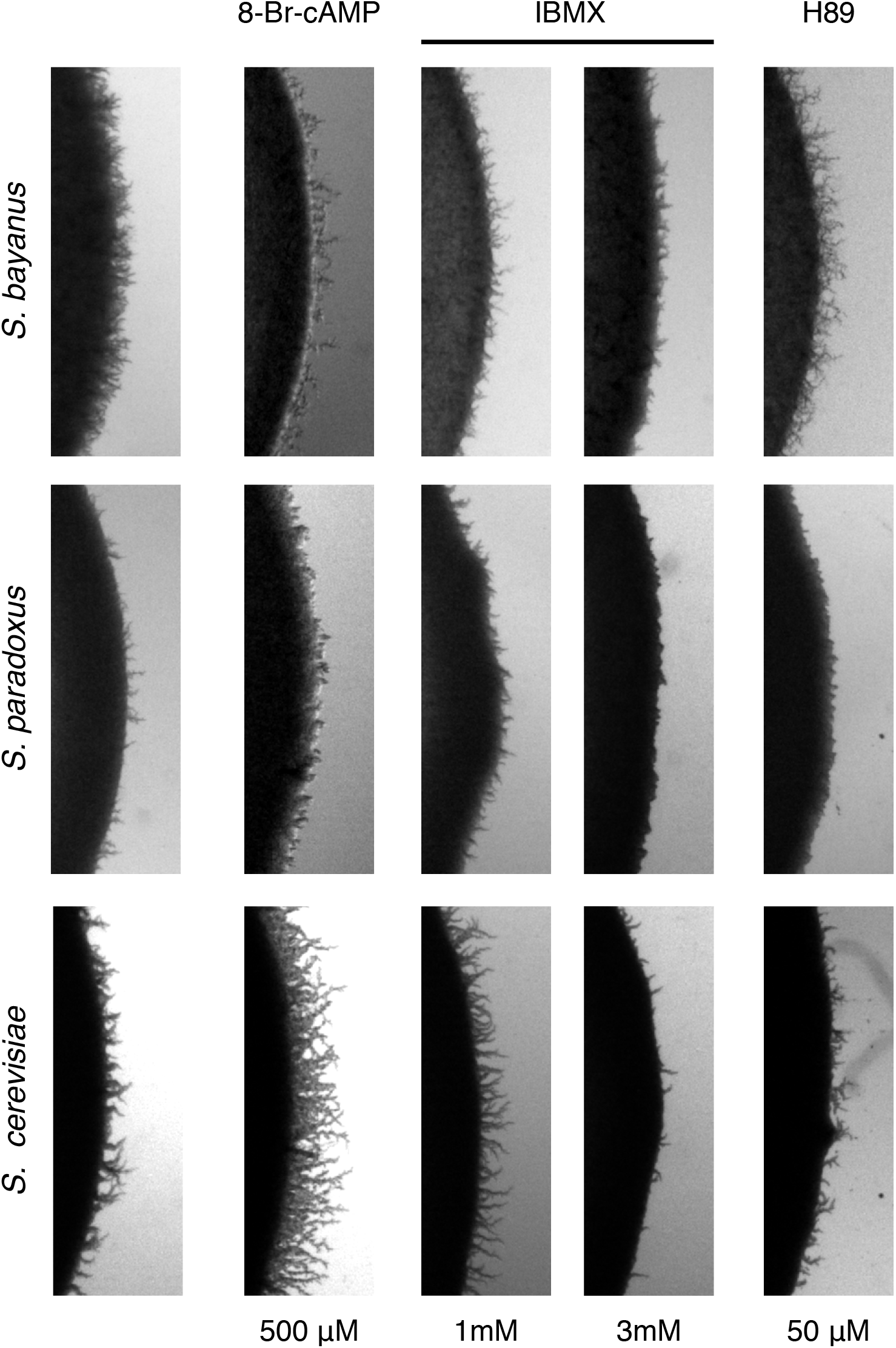
Pharmacological modulations of cAMP levels. Intracellular cAMP levels were modified using indicated drugs in three *Saccharomyces* species. Effects of 8-Br-cAMP were similar to exogenous cAMP treatment. Low concentrations of IBMX reflect the effects of increased intracellular cAMP levels but a higher concentration blocks the response likely due to collapse of the cAMP signaling. PKA inhibitor H89 had a strong effect on *S. paradoxus*.

**Figure S5.**
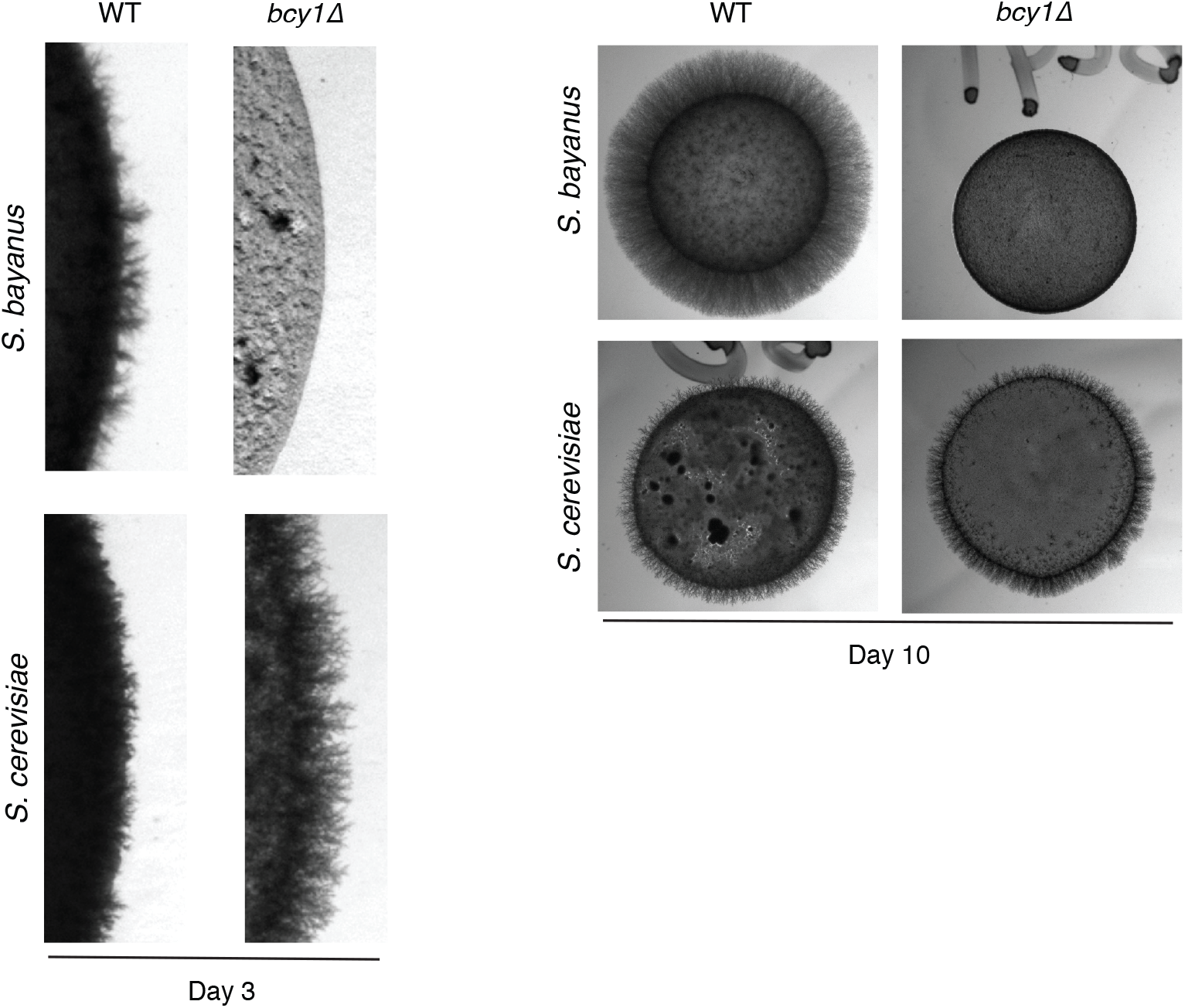
Hyperactivation of PKA and divergent pseudohyphal responses, *bcy1* at day 3 and 10. The *bcy1* mutation causes a complete loss of pseudohyphal differentiation in *S. bayanus*, consistent with effects of increased cAMP levels. The bcy1 mutant exhibits hyperfilamentation in *S. cerevisiae*.

**Figure S6.**
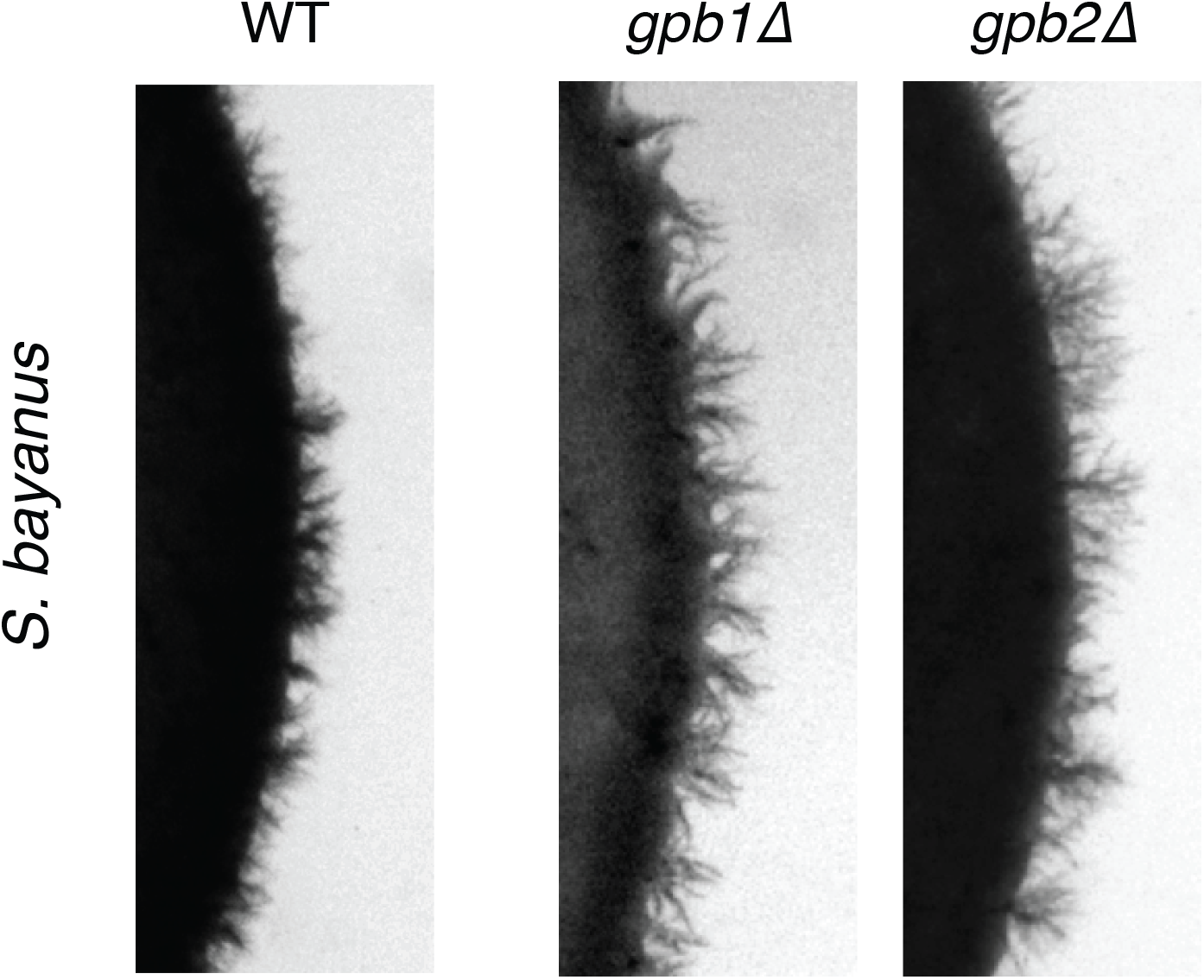
Deletions of GPB1 and GPB2 lead to a slight increase in filamentous growth in *S. bayanus*. The change is comparable to reported observations for the same deletion mutants in *S. cerevisiae*.

## Supplementary Tables

**Table S1.** *S. cerevisiae* strains assayed for pseudohyphal growth ability (0 = no pseudohyphal growth, 1 = pseudohyphal) on SLAD plates, and in the presence of exogenous cAMP added to the growth medium.

| Strain Number | Strain Name | Origin | Psh | 1 mM<br>cAMP | 3 mM<br>cAMP | 10 mM<br>cAMP |
| --- | --- | --- | --- | --- | --- | --- |
| PMY 011 | YPS602 | Oak | 0 | 1 | 1 | 1 |
| PMY 012 | YPS606 | Oak | 0 | 1 | 1 | 1 |
| PMY 014 | YPS623 | Oak | 0 | 1 | 1 | 1 |
| PMY 015 | YPS630 | Oak | 0 | 1 | 1 | 1 |
| PMY 017 | YPS670 | Oak | 0 | 1 | 1 | 1 |
| PMY 018 | YPS681 | Oak | 0 | 1 | 1 | 1 |
| PMY 070 | EM93 | Fig | 1 | 1 | 1 | 1 |
| PMY 072 | PMY072 | Vineyard | 1 | 1 | 1 | 1 |
| PMY 074 | PMY074 | Vineyard | 1 | 1 | 1 | 1 |
| PMY 083 | PMY083 | Vineyard | 1 | 1 | 1 | 1 |
| PMY 084 | PMY084 | Vineyard | 1 | 1 | 1 | 1 |
| PMY 086 | PMY086 | Vineyard | 0 | 1 | 1 | 1 |
| PMY 087 | PMY087 | Vineyard | 0 | 1 | 1 | 1 |
| PMY 088 | PMY088 | Vineyard | 0 | 1 | 1 | 1 |
| PMY 093 | PMY093 | Vineyard | 0 | 1 | 1 | 1 |
| PMY 094 | PMY094 | Vineyard | 0 | 1 | 1 | 1 |
| PMY 095 | PMY095 | Vineyard | 0 | 1 | 1 | 1 |
| PMY 110 | PMY110 | Vineyard | 1 | 1 | 1 | 1 |
| PMY 111 | PMY111 | Vineyard | 1 | 1 | 1 | 1 |
| PMY 112 | PMY112 | Vineyard | 1 | 1 | 1 | 1 |
| PMY 113 | YJM336 | Clinical | 1 | 1 | 1 | 1 |
| PMY 116 | YJM431 | Laboratory | 0 | 0 | 1 | 1 |
| PMY 119 | YJM454 | Clinical | 1 | 1 | 1 | 1 |
| PMY 120 | F4852 | Clinical | 1 | 1 | 1 | 1 |
| PMY 121 | Y55 | Laboratory | 1 | 1 | 1 | 1 |
| PMY 123 | SK1 | Laboratory | 1 | 1 | 1 | 1 |
| PMY 127 | YJM128 | Clinical | 1 | 1 | 1 | 1 |
| PMY 128 | YJM128 | Clinical | 1 | 1 | 1 | 1 |
| PMY 129 | CBS1227 | Clinical | 1 | 1 | 1 | 1 |
| PMY 131 | YJM222 | Clinical | 1 | 1 | 1 | 1 |
| PMY 133 | YJM224 | Distillery | 1 | 1 | 1 | 1 |
| PMY 137 | VMC132B | Clinical | 1 | 1 | 1 | 1 |
| PMY 140 | YJM277 | Clinical | 1 | 1 | 1 | 1 |
| PMY 141 | 90-59 | Clinical | 1 | 1 | 1 | 1 |
| PMY 144 | YJM311 | Clinical | 1 | 1 | 1 | 1 |
| PMY 147 | YJM334 | Vineyard | 1 | 1 | 1 | 1 |

**Table S2.**
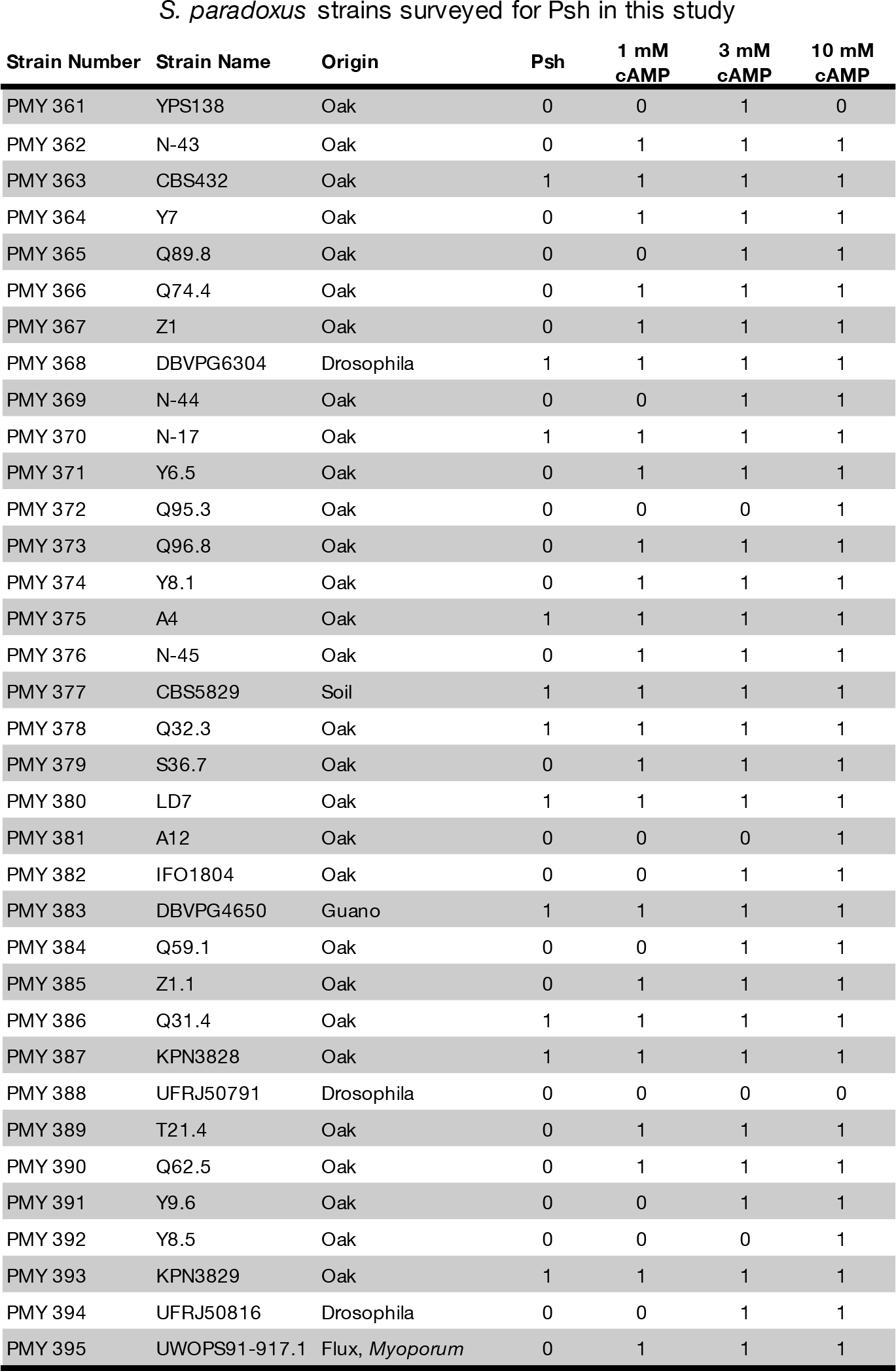
*S. paradoxus* strains assayed for pseudohyphal growth ability (0 = no pseudohyphal growth, 1 = pseudohyphal) on SLAD plates, and in the presence of exogenous cAMP added to the growth medium.

**Table S3.** *S. bayanus* strains assayed for pseudohyphal growth ability (0 = no pseudohyphal growth, 1 = pseudohyphal) on SLAD plates, and in the presence of exogenous cAMP added to the growth medium.

| Strain Number | Strain Name | Origin | Psh | 1mM<br>cAMP | 3mM<br>cAMP | 10mM<br>cAMP |
| --- | --- | --- | --- | --- | --- | --- |
| PMY 640 | NCYC365 | Apple juice | 1 | 1 | 0 | 0 |
| PMY 641 | NCYC2578 | Turbid Beer | 1 | 1 | 0 | 0 |
| PMY 642 | CBS7001 | Mesophylax adopersus (insect) | 1 | 1 | 0 | 0 |
| PMY 643 | YJM520 | Fermenting juice of Taffert apples | 1 | 1 | 0 | 0 |
| PMY 644 | YJM519 | Pear Juice | 0 | 0 | 0 | 0 |
| PMY 645 | NCYC509 | Juice of Ribes nigrum (Blackcurrant) | 1 | 1 | 0 | 0 |
| PMY 660 | GL 222 | Unknown | 1 | 1 | 0 | 0 |
| PMY 661 | GL 274 | Unknown | 0 | 0 | 0 | 0 |
| PMY 668 | VKM Y-361 | Czech Wine | 0 | 0 | 0 | 0 |
| PMY 669 | NRRL Y-969 | Unknown | 1 | 1 | 0 | 0 |
| PMY 670 | VKM Y-1146 | Grape berries | 1 | 1 | 0 | 0 |
| PMY 734 | NBRC539 | Unknown | 1 | 1 | 0 | 0 |
| PMY 735 | NCYC114 | Unknown | 1 | 1 | 0 | 0 |
| PMY 736 | NBRC10558 | Must of soft fruit | 0 | 0 | 0 | 0 |
| PMY 911 | ZP555 | Oak | 1 | 0 | 0 | 0 |
| PMY 912 | ZP556 | Oak | 0 | 0 | 0 | 0 |
| PMY 937 | NCYC686 | Spoiled Coca-Cola | 1 | 1 | 0 | 0 |
| PMY 938 | NCYC762 | Palm Wine | 0 | 0 | 0 | 0 |
| PMY 939 | NCYC1322 | Irish Brewery | 0 | 0 | 0 | 0 |
| PMY 940 | NCYC1323 | Australian Brewery | 1 | 0 | 0 | 0 |
| PMY 941 | NCYC1324 | Scottish Brewery | 0 | 0 | 0 | 0 |
| PMY 942 | NCYC1341 | British Brewery | 0 | 0 | 0 | 0 |
| PMY 943 | NCYC3066 | Fermenting Saurkraut | 0 | 0 | 0 | 0 |
| PMY 944 | NCYC3359 | Cider apples | 1 | 1 | 0 | 0 |
| PMY 947 | NCYC1326 | British Brewery | 0 | 0 | 0 | 0 |
| PMY 948 | NCYC1342 | British Brewery | 1 | 1 | 1 | 1 |
| PMY 949 | NCYC965 | British Brewery | 1 | 1 | 1 | 1 |
| PMY 952 | NCYC966 | British Brewery | 1 | 1 | 1 | 1 |
| PMY 953 | NCYC967 | British Brewery | 1 | 1 | 1 | 1 |
| PMY 955 | NCYC969 | British Brewery | 0 | 0 | 0 | 0 |
| PMY 956 | NCYC984 | European Brewery | 0 | 0 | 0 | 0 |
| PMY 957 | NCYC985 | European Brewery | 1 | 1 | 1 | 1 |
| PMY 958 | NCYC986 | European Brewery | 1 | 1 | 1 | 1 |
| PMY 959 | NCYC987 | European Brewery | 1 | 1 | 1 | 1 |
| PMY 969 | NCYC988 | European Brewery | 0 | 0 | 0 | 0 |
| PMY 970 | NCYC989 | European Brewery | 1 | 1 | 1 | 1 |

**Table S4.** Summary of the effects of exogenous cAMP treatment on pseudohyphal growth for strains of *S. cerevisiae, S. paradoxus*, and *S. bayanus*.

| S. bayanus cAMP treatment |  |  |
| --- | --- | --- |
|  | Pseudohyphal | Non-pseudohyphal |
| Decrease | 12 | N/A |
| Increase | 0 | 0 |
| No-observable Change | 10 | 14 |

| S. paradoxus cAMP treatment |  |  |
| --- | --- | --- |
|  | Pseudohyphal | Non-pseudohyphal |
| Decrease | 0 | N/A |
| Increase | 11 | 24 |
| No-observable Change | 0 | 0 |

| S. cerevisiae cAMP treatment |  |  |
| --- | --- | --- |
|  | Pseudohyphal | Non-pseudohyphal |
| Decrease | 0 | N/A |
| Increase | 23 | 13 |
| No-observable Change | 0 | 0 |

**Table S5.** The effects on pseudohyphal growth of null mutations of genes involved in cAMP-PKA signaling in *S. cerevisiae* and *S. bayanus*. The pseudohyphal response in mutant strains was evaluated relative to the wild type phenotype in the same genetic background after 72 hours of growth on SLAD medium. Phenotypes were classified as increasing (+), decreasing (−), no change (∅), no change or slight increase (∅/+), and no-change or slight decrease (∅/−).

| Category | Gene | *S. cerevisiae* | *S. bayanus* |
| --- | --- | --- | --- |
| 1 | *gpa2Δ* | - | - |
| 1 | *pde2Δ* | ∅ | ∅ |
| 1 | *phd1Δ* | - | - |
| 1 | *sfl1Δ* | + | + |
| 1 | *tpk1Δ* | + | + |
| 1 | *tpk2Δ* | - | - |
| 1 | *tpk3Δ* | + | + |
| 1 | *gpb1Δ* | + | ∅/+ |
| 1 | *gpb2Δ* | + | ∅/+ |
| 1 | *ras1Δ* | ∅ | ∅/- |
| 1 | *ras2Δ* | - | ∅/- |
| 2 | *bcy1Δ* | + | - |
| 2 | *flo8Δ* | - | + |
| 2 | *flo11Δ* | - | ∅ |
| 2 | *ira2Δ* | + | - |
| 2 | *pde1Δ* | + | - |

**Table S6.** Protein sequence alignment for key components of the cAMP-PKA signaling network indicates a high degree of conservation across the *Saccharomyces sensu stricto* clade.

|  | % ID / Similarity<br>S. paradoxus | % ID / Similarity<br>S. uvarum | % ID / Similarity<br>S. bayanus | Comparison<br>Matrix |
| --- | --- | --- | --- | --- |
| **Bcy1** | 98.3 / 98.3 | 93.8 / 96.6 | 93.8 / 96.6 | BLOSUM 62 |
| **Cyr1** | 95.1 / 97.1 | 89.4 / 93.9 | 89.4 / 93.9 | BLOSUM 62 |
| **Flo8** | 87.0 / 90.6 | NA | 66.7 / 78.3 | BLOSUM 62 |
| **Gbp2** | 72.4 / 76.8 | 75.8 / 85.7 | 75.8 / 85.7 | BLOSUM 62 |
| **Gpa2** | 92.7 / 95.4 | 79.6 / 87.2 | 79.6 / 87.2 | BLOSUM 62 |
| **Gpb1** | 91.1 / 95.0 | 79.4 / 88.9 | 79.4 / 88.9 | BLOSUM 62 |
| **Ira2** | 95.5 / 98.0 | NA | 87.3 / 94.4 | BLOSUM 62 |
| **Pde1** | 95.9 / 98.4 | 84.9 / 92.2 | 84.9 / 92.2 | BLOSUM 62 |
| **Pde2** | 95.1 / 99.0 | 84.3 / 91.7 | 84.3 / 91.7 | BLOSUM 62 |
| **Phd1** | 91.3 / 94.8 | 71.2 / 81.7 | 69.5 / 80.1 | BLOSUM 62 |
| **Ras2** | 93.2 / 96.0 | 83.0 / 88.2 | 83.0 / 88.2 | BLOSUM 62 |
| **Sfl1** | 92.2 / 94.7 | 65.4 / 71.4 | 75.9 / 83.9 | BLOSUM 62 |
| **Tpk1** | 97.0 / 97.5 | 91.7 / 95.0 | 91.7 / 95.0 | BLOSUM 62 |
| **Tpk2** | 97.9 / 99.2 | 82.8 / 84.2 | 93.5 / 95.1 | BLOSUM 62 |
| **Tpk3** | 96.7 / 98.0 | 93.0 / 95.5 | 91.7 / 94.2 | BLOSUM 62 |

**Table S7.** List of mutant strains used in this study.

| Species | Strain | Genotype | Reference |
| --- | --- | --- | --- |
| ***S. bayanus*** | PMY 0640 | NCYC365 | Ludo & McCusker, 1999 |
|  | PMY 1392 | *gpa2Δ:: kanMX* | This study |
|  | PMY 1394 | *ste7Δ:: kanMX* | This study |
|  | PMY 1396 | *flo8Δ:: kanMX* | This study |
|  | PMY 1442 | *dig1Δ:: kanMX* | This study |
|  | PMY 1445 | *ras2Δ:: kanMX* | This study |
|  | PMY 1446 | *tec1Δ:: kanMX* | This study |
|  | PMY 1448 | *tpk1Δ:: kanMX* | This study |
|  | PMY 1450 | *ste12Δ:: kanMX* | This study |
|  | PMY 1455 | *pde1Δ:: kanMX* | This study |
|  | PMY 1459 | *tpk3Δ:: kanMX* | This study |
|  | PMY 1460 | *sch9Δ:: kanMX* | This study |
|  | PMY 1461 | *pde2Δ:: kanMX* | This study |
|  | PMY 1462 | *bcy1Δ:: kanMX* | This study |
|  | PMY 1479 | *tpk2Δ:: kanMX* | This study |
|  | PMY 1480 | *rim15Δ:: kanMX* | This study |
|  | PMY 1481 | *ras1Δ:: kanMX* | This study |
|  | PMY 1482 | *gpb1Δ:: kanMX* | This study |
|  | PMY 1483 | *gpb2Δ:: kanMX* | This study |
|  | PMY 1484 | *ira2Δ:: kanMX* | This study |
|  | PMY 1596 | *msn2Δ:: kanMX* | This study |
|  | PMY 1600 | *flo11Δ:: kanMX* | This study |
|  | PMY 1603 | *phd1Δ:: kanMX* | This study |
|  | PMY 1606 | *sfl1Δ:: kanMX* | This study |
| ***S. cerevisiae*** | PMY 0485 | *gpa2Δ:: kanMX* | Lorenz & Heitman, 1997 |
|  | PMY 0487 | *ste12Δ:: kanMX* | Lorenz & Heitman, 1997 |
|  | PMY 0491 | *flo8Δ:: kanMX* | Liu, *et.al.*, 1996 |
|  | PMY 0498 | *ira2Δ:: kanMX* | Drees, *et.al.*, 2005 |
|  | PMY 0499 | *ras2Δ:: kanMX* | Drees, *et.al.*, 2005 |
|  | PMY 0501 | *tec1Δ:: kanMX* | Drees, *et.al.*, 2005 |
|  | PMY 0506 | *tpk1Δ:: kanMX* | Drees, *et.al.*, 2005 |
|  | PMY 0507 | *tpk2Δ:: kanMX* | Drees, *et.al.*, 2005 |
|  | PMY 0508 | *tpk3Δ:: kanMX* | Drees, *et.al.*, 2005 |
|  | PMY 0746 | *pde1Δ:: kanMX* | Drees, *et.al.*, 2005 |
|  | PMY 0748 | *flo11Δ:: kanMX* | Drees, *et.al.*, 2005 |
|  | PMY 0750 | *pde2Δ:: kanMX* | Drees, *et.al.*, 2005 |
|  | PMY 0827 | *ste7Δ:: kanMX* | Lorenz & Heitman, 1997 |
|  | PMY 0843 | *phd1Δ:: kanMX* | Lorenz & Heitman, 1998 |
|  | PMY 0861 | *bcy1Δ:: kanMX* | Pan & Heitman, 1999 |
|  | PMY 0982 | *rim15Δ:: kanMX* | Pan & Heitman, 1999 |
|  | PMY 1039 | *msn2Δ:: kanMX* | Pan & Heitman, 1999 |
|  | PMY 1048 | *sch9Δ:: kanMX* | Lorenz, *et.al.*, 1999 |
|  | PMY 1064 | *sfl1Δ:: kanMX* | Pan & Heitman, 2002 |

**Table S8.**
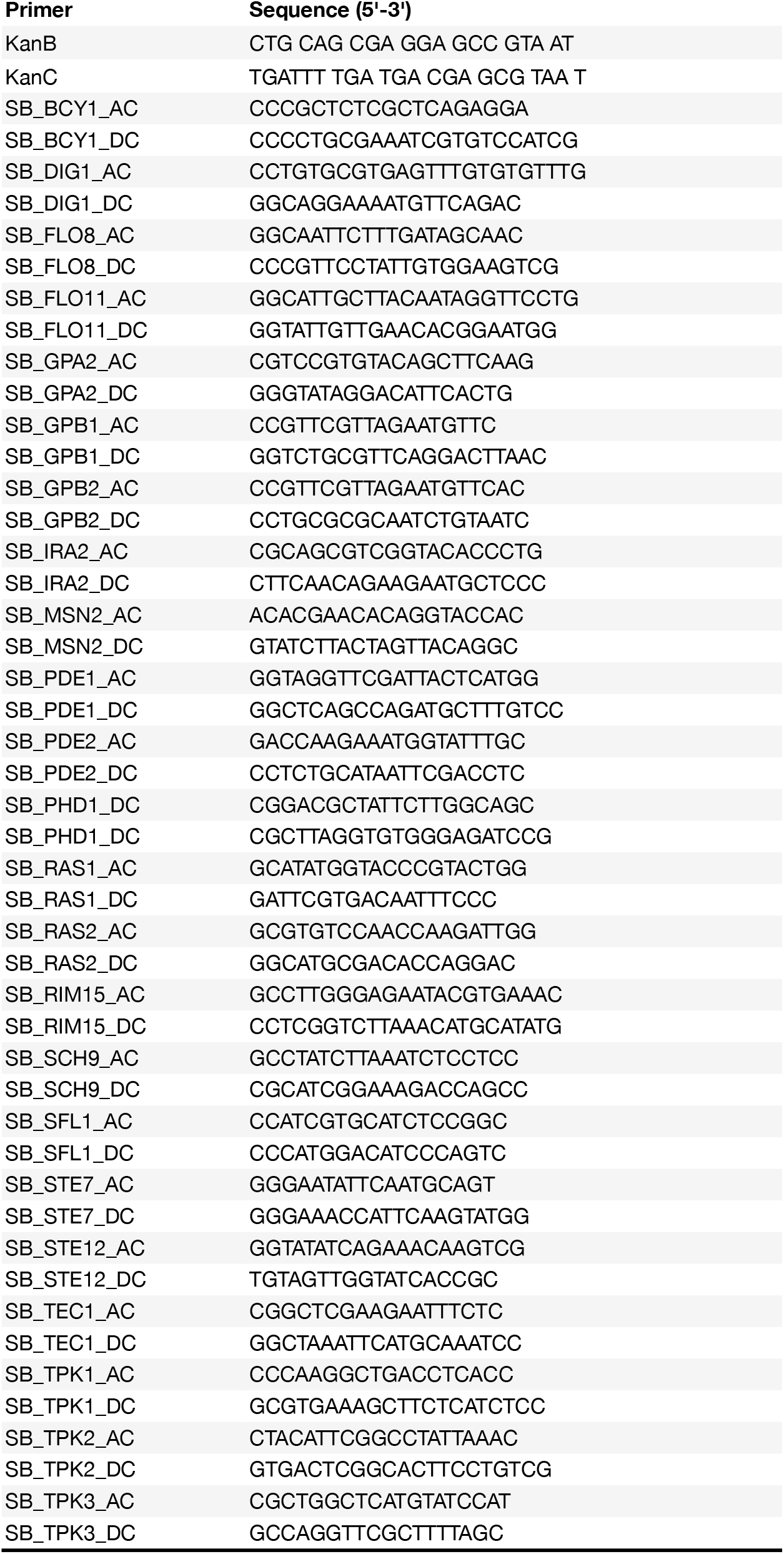
List of primers that were used for confirmation-PCR to validate deletion of the targeted locus for each null mutant.

